# Plant-derived Cellulose Scaffolds for Bone Tissue Engineering

**DOI:** 10.1101/2020.01.15.906677

**Authors:** Maxime Leblanc Latour, Maryam Tarar, Ryan J. Hickey, Charles M. Cuerrier, Isabelle Catelas, Andrew E. Pelling

**Affiliations:** Department of Physics, University of Ottawa, 150 Louis-Pasteur, Ottawa, ON, K1N 6N5, Canada; Department of Biology, University of Ottawa, Gendron Hall, 30 Marie Curie, Ottawa, ON, K1N 6N5, Canada; Department of Mechanical Engineering, University of Ottawa, 161 Louis Pasteur, Ottawa, ON, K1N 6N5, Canada; Department of Surgery, The Ottawa Hospital – General Campus, 501 Smyth Road, Ottawa, ON, K1H 8L6, Canada; Department of Biochemistry, Microbiology and Immunology, University of Ottawa, 451 Smyth Road, Ottawa, ON, K1H 8M5, Canada; Institute for Science, Society and Policy, University of Ottawa, Desmarais Building, 55 Laurier Ave. East, Ottawa, ON, K1N 6N5, Canada; SymbioticA, School of Human Sciences, University of Western Australia, 35 Stirling Highway, Perth, Western Australia, 6009, Australia

## Abstract

Plant-derived cellulose biomaterials have recently been utilized in several tissue engineering applications. Naturally-derived cellulose scaffolds have been shown to be highly biocompatible *in vivo*, possess structural features of relevance to several tissues, as well as support mammalian cell invasion and proliferation. Recent work utilizing decellularized apple hypanthium tissue has shown that it possesses a pore size and properties similar to trabecular bone. In the present study, we examined the potential of apple-derived cellulose scaffolds for bone tissue engineering (BTE). Confocal microscopy revealed that the scaffolds had a suitable pore size for BTE applications. To analyze their *in vitro* mineralization potential, MC3T3-E1 pre-osteoblasts were seeded in either bare cellulose scaffolds or in composite scaffolds composed of cellulose and collagen I. Following chemically-induced differentiation, scaffolds were mechanically tested and evaluated for mineralization. The Young’s modulus of both types of scaffolds significantly increased after cell differentiation. Alkaline phosphatase and Alizarin Red staining further highlighted the osteogenic potential of the scaffolds. Histological sectioning of the constructs revealed complete invasion by the cells and mineralization throughout the entire constructs. Finally, scanning electron microscopy demonstrated the presence of mineral aggregates deposited on the scaffolds after differentiation, and energy-dispersive spectroscopy confirmed the presence of phosphate and calcium. In summary, our results indicate that plant-derived cellulose is a promising scaffold candidate for bone tissue engineering applications.

## Introduction

Large bone defects caused by injury or disease often require biomaterial grafts to completely regenerate [1]. Current techniques designed to enhance bone tissue regeneration commonly employ autologous, allogeneic, xenogeneic, or synthetic grafts [2]. Autologous bone grafting, in which the material is derived from the patient, is considered the “gold standard” grafting practice in large bone defect repairs, but there are several drawbacks including size and shape limitations, tissue availability, and donor site morbidity [3]. In addition, autologous grafting procedures are prone to infections, subsequent fractures, hematoma formation at the donor or repaired site, and post-operative pain [4]. Bone tissue engineering (BTE) provides a potential alternative to traditional bone grafting methods [5]. It combines the use of structural biomaterials and cells to create new functional bone tissue. The biomaterials used for BTE require similar architecture and mechanical properties to the native bone matrix [6]. Previous studies have shown that the optimal pore size for biomaterials used for BTE is approximately 100-200 μm [7], and elastic modulus is 0.1 to 20 GPa depending on the grafting site [8]. Moreover, the porosity and pore interconnectivity are two important factors that affect cell migration, nutrient diffusion, and angiogenesis [8]. BTE has shown promising results with a diverse set of biomaterials developed as an alternative to bone grafts. These biomaterials include osteoinductive materials, hybrid materials, and advanced hydrogels [8]. Osteoinductive materials induce the formation of *de novo* bone structure. Hybrid materials are made of synthetic and/or natural polymers [8]. Advanced hydrogels mimic the extracellular matrix (ECM) and deliver the required bioactive agents to promote bone tissue integration [8]. Hydroxyapatite is a traditional material choice for BTE due to its biocompatibility and composition [9]. Another type of biomaterial for BTE is bioactive glass, which stimulates specific cell responses to activate genes necessary for osteogenesis [10], [11]. Biodegradable polymers such as poly(glycolic acid) and poly(lactic acid) are also widely used for BTE [12]. Finally, natural (or naturally-derived) polymers such as chitosan, chitin, and bacterial cellulose have also shown promising results for BTE [13].

Although these polymers, either natural or synthetic, show some potential in BTE, extensive protocols are usually required to obtain a functional biomaterial with a desired macrostructure. Conversely, native macroscopic cellulose structures can be derived from various plants. In a previous study, our group demonstrated that cellulose-based scaffolds derived from plants, using a simple surfactant treatment, can be used as a material for various tissue reconstruction by taking advantage of the native structure of the plant [14]. Furthermore, these biomaterials can be used for *in vitro* mammalian cell culture [14], are biocompatible, and can become spontaneously vascularized subcutaneously [14]–[16]. Our group and others have shown that these biomaterials can be sourced from specific plants according to the intended application [14]–[18]. For instance, the vascular structure from plant stems and leaves displays a similar structure as the one found in animal tissue [18]. Plant-derived cellulose scaffolds can also easily be carved into specific shapes and treated to alter their surface biochemistry [16]. In a recent study, we included a salt buffer in the decellularization process, which resulted in an increase in cell attachment, both *in vitro* and *in vivo* [16]. In the same study, we showed that plant-derived cellulose can be used in composite biomaterials by casting hydrogels onto the scaffold surface. Finally, it has been shown that apple hypanthium tissue can provide a bone-like architecture, with interconnected pores ranging from 100 to 200 μm in diameter [14]. In the present proof-of-concept study, we demonstrate that apple-derived cellulose scaffolds can act as a suitable biomaterial for BTE. While other studies have shown promising results using bacterial cellulose for BTE [19], plant-derived cellulose has not been previously employed for this particular application. Importantly, hypanthium tissue possesses a microstructure with geometric characteristics similar to trabecular bone [7].

Scaffolds derived from apple hypanthium tissue were prepared in two formulations that we previously described [14]–[16]. MC3T3-E1 pre-osteoblasts were seeded on bare cellulose scaffolds or composite scaffolds composed of a collagen hydrogel embedded in cellulose scaffolds [16]. Both scaffold preparations supported cellular invasion and proliferation at which point the scaffolds were placed in osteoinductive medium. After cell osteogenic differentiation, both scaffold types depicted a higher Young’s modulus, alkaline phosphatase activity, as well as calcium deposition and mineralization. Results confirmed the potential of these low cost, sustainable, and renewable plant-derived cellulose scaffolds for BTE applications.

## Materials and Methods

### Scaffold preparation

Samples were prepared following established methods [16]. Briefly, McIntosh apples (Canada Fancy) were cut in 8 mm-thick slices with a mandolin slider. The hypanthium tissue of the apple slices was cut into squares of 5 mm by 5 mm. The square tissues were decellularized in 0.1% sodium dodecyl sulfate (SDS, Fisher Scientific, Fair Lawn, NJ) for two days. Decellularized samples were then washed in deionized water, followed by an overnight incubation in 100 mM CaCl_2_ to remove the remaining surfactant. The samples were subsequently sterilized with 70% ethanol for 30 min, washed with deionized water, and placed in a 24-well culture plate prior to cell seeding. The scaffolds (8-mm thick) were either left untreated (bare scaffolds) or coated with a collagen solution (composite hydrogel scaffolds), as explained below.

MC3T3-E1 Subclone 4 cells (ATCC® CRL-2593™, Manassas, VA) were maintained at 37°C in a humidified atmosphere of 95% air and 5% CO_2_. The cells were cultured in Minimum Essential Medium (α-MEM, ThermoFisher, Waltham, MA), supplemented with 10% fetal bovine serum (FBS Hyclone Laboratories Inc., Logan, UT) and 1% penicillin/streptomycin (Hyclone Laboratories Inc.), and were allowed to grow to 80% confluency before being trypsinized. They were then resuspended at 10^5^ cells/mL in either α-MEM or a 1.5 g/L collagen solution, as follows, for the preparation of the bare scaffolds or the scaffolds coated with a collagen solution, respectively. Briefly, the collagen solution was prepared by mixing 50% (v/v) of 3 mg/mL type 1 collagen (ThermoFisher) with 2.5% of 1N NaOH, 1% of FBS, 10% of 10x phosphate-buffered saline (PBS, ThermoFisher), and 36.5% of sterile deionized water at 4°C. A 40 μL aliquot of cell suspension, in either α-MEM or a 1.5 g/L collagen solution, was pipetted on the scaffolds. The cells were left to adhere for 1h in cell culture conditions (i.e., at 37°C in a humidified atmosphere of 95% air and 5% CO_2_). Subsequently, 2 mL of culture medium were added to each culture well. Culture medium was changed every 2-3 days, for 14 days. After these 14 days of incubation, differentiation of MC3T3-E1 cells was induced by adding 50 μg/mL ascorbic acid and 4 mM sodium phosphate to the culture medium (differentiation medium). Differentiation medium was changed every 3-4 days, for 4 weeks. Scaffolds in non-differentiation culture medium (without the supplements to induce differentiation) were incubated for the same period of time, with the same medium change frequency, and served as a negative control. All subsequent analyses were conducted at the end of this 4-week incubation period. Finally, the decellularized apple scaffolds as well as the cell-seeded bare and composite scaffolds were imaged after the 4-week incubation using a 12 megapixel digital camera.

### Pore size measurements and cell distribution analysis using confocal laser scanning microscopy

To measure the scaffold pore size, decellularized apple scaffolds (prior to collagen treatment and MC3T3-E1 cell seeding) were thoroughly washed with PBS and stained with 1 mL of 10% (v/v) Calcofluor White solution (Sigma-Aldrich, St. Louis, MO) for 25 min in the dark and at room temperature. Subsequently, scaffolds (n=3) were washed with PBS and were imaged with a high-speed resonant confocal laser scanning microscope (Nikon Ti-E A1-R; Nikon, Mississauga, ON). ImageJ software [20] was used to process and analyze the confocal images. Briefly, maximum projections in the Z axis were created and the Find Edges function was used to highlight the edge of the pores. A total of 54 pores were analyzed (6 pores in 3 randomly selected area per scaffold, with n=3 scaffolds). Pores were manually traced using the freehand selection tool in ImageJ. The selections were fit as an ellipse to output the major axis length.

To analyze MC3T3-E1 cell distribution in the scaffolds, bare and composite cell-seeded scaffolds (n=3 for each experimental condition) were thoroughly washed with PBS and fixed with 4% paraformaldehyde for 10 min. They were then extensively washed with deionized water before permeabilizing the cells with a Triton-X 100 solution (ThermoFisher) for 5 min, and washed again with PBS. Staining of the scaffolds was carried out as previously described [14], [16]. Briefly, the scaffolds were incubated in 1% periodic acid (Sigma-Aldrich) for 40 min. After rinsing with deionized water, they were incubated in 100 mM sodium metabisulphite (Sigma-Aldrich) and 0.15 M hydrochloric acid (ThermoFisher), supplemented with 100 μg/mL propidium iodide (Invitrogen, Carlsbad, CA) for 2h in the dark and at room temperature. Finally, they were washed in PBS, stained with 5 mg/mL DAPI (ThermoFisher) for 10 min in the dark, washed again, and stored in PBS prior to imaging. The cell-seeded surfaces of the scaffolds were imaged with a high-speed resonant confocal laser scanning microscope (Nikon Ti-E A1-R). ImageJ software [20] was used to process the confocal images and create a maximum projection in the Z axis for image analysis.

### Young’s modulus measurements

Young’s modulus measurements of the scaffolds (n=3 for each experimental condition) with non-differentiated and differentiated cells were obtained using a custom-built uniaxial compression apparatus. Decellularized apple-derived cellulose scaffolds without cells were used as a control. The force and position was recorded with a 150g load cell (Honeywell) and an optical ruler. The force-displacement curves were obtained by compressing the samples at a constant rate of 3 mm min^−1^ and a maximum strain of 10%. The Young’s modulus was obtained by fitting the linear portion of the stress-strain curve.

### Alkaline phosphatase and Alizarin Red S staining

Before staining with either 5-bromo-4-chloro-3’-indolyphosphate and nitro-blue tetrazolium (BCIP/NBT, ThermoFisher) or Alizarin Red S (ARS, Sigma-Aldrich), scaffolds were washed three times with PBS (without Ca^2+^ and Mg^2+^, Hyclone Laboratories Inc.) and fixed with 10% neutral buffered formalin for 30 min.

BCIP/NBT was used to assess the alkaline phosphatase (ALP) activity of cell-seeded scaffolds. BCIP/NBT staining solution was prepared by dissolving one BCIP/NBT tablet (Sigma-Aldrich) in 10 mL of deionized water. After fixation, the scaffolds (n=3 for each experimental condition) were washed with a 0.05% Tween solution and stained with BCIP/NBT for 20 min at room temperature. Finally, they were washed with 0.05% Tween and stored in PBS (without Ca^2+^ and Mg^2+^) prior to imaging.

ARS was used to assess calcium deposition and mineralization of the scaffolds. After fixation, the scaffolds (n=3 for each experimental condition) were washed with deionized water and exposed to 2% (w/v) ARS for 1h at room temperature. They were then washed with deionized water to remove the excess ARS staining solution and stored in PBS (without Ca^2+^ and Mg^2+^) prior to imaging.

Finally, all scaffolds were imaged using a 12 megapixel digital camera.

### Histological analysis

Scaffolds (n=1 for scaffolds in non-differentiation medium and n=2 for scaffolds in differentiation medium) were fixed in 4% para-formaldehyde for 48h, and stored in 70% ethanol before paraffin embedding. Embedding, sectioning, and staining were performed by the PALM Histology Core Facility of the University of Ottawa. Briefly, 5 μm-thick serial sections were stained with hematoxylin and eosin (H&E; ThermoFisher) or Von Kossa (VK; ThermoFisher), starting 1 mm inside the scaffolds. Slides (n=2 per scaffold) were imaged using a Zeiss AXIOVERT 40 CFL microscope (Zeiss, Toronto, ON) to evaluate cell infiltration (H&E) and mineralization (VK). Image analysis was performed using ImageJ software.

### Mineralization analysis using scanning electron microscopy and energy-dispersive spectroscopy

Scaffolds (n=3 for each experimental condition) were fixed in 4% para-formaldehyde for 48h, followed by serial dehydration in increasing concentrations of ethanol (from 50% to 100%), as previously described [21]. Samples where then dried using a critical point dryer. Dried samples were gold-coated to a final coating thickness of 5 nm. Scanning electron microscopy (SEM) images were acquired with a JEOL JSM-7500F FESEM scanning electron microscope (JEOL, Peabody, MA) at 2 kV. Energy-dispersive spectroscopy (EDS) was performed on bare scaffolds and composite hydrogel scaffolds seeded with MC3T3-E1 cells. Three different areas of each scaffold surface were analyzed for mineral aggregates.

### Statistical analysis

Young’s modulus data are reported as mean ± standard error of the mean (S.E.M.). The data were assumed to be normally distributed. Statistical analysis was performed using a one-way ANOVA followed by Tukey post-hoc tests. A value of *p* < 0.05 was considered to be statistically significant.

## Results

### Scaffold imaging and pore size measurements

Complete removal of native cellular components of the apple tissue was achieved after SDS and CaCl_2_ treatments (Figure 1A). White calcium deposits were observed throughout the bare and composite hydrogel scaffolds cultured with differentiation medium for 4 weeks (Figure 1B and C, respectively). Both types of scaffolds with differentiated cells had a distinct opaque white colour that was absent in the control scaffolds without cells (Figure 1A).

**Figure 1:**
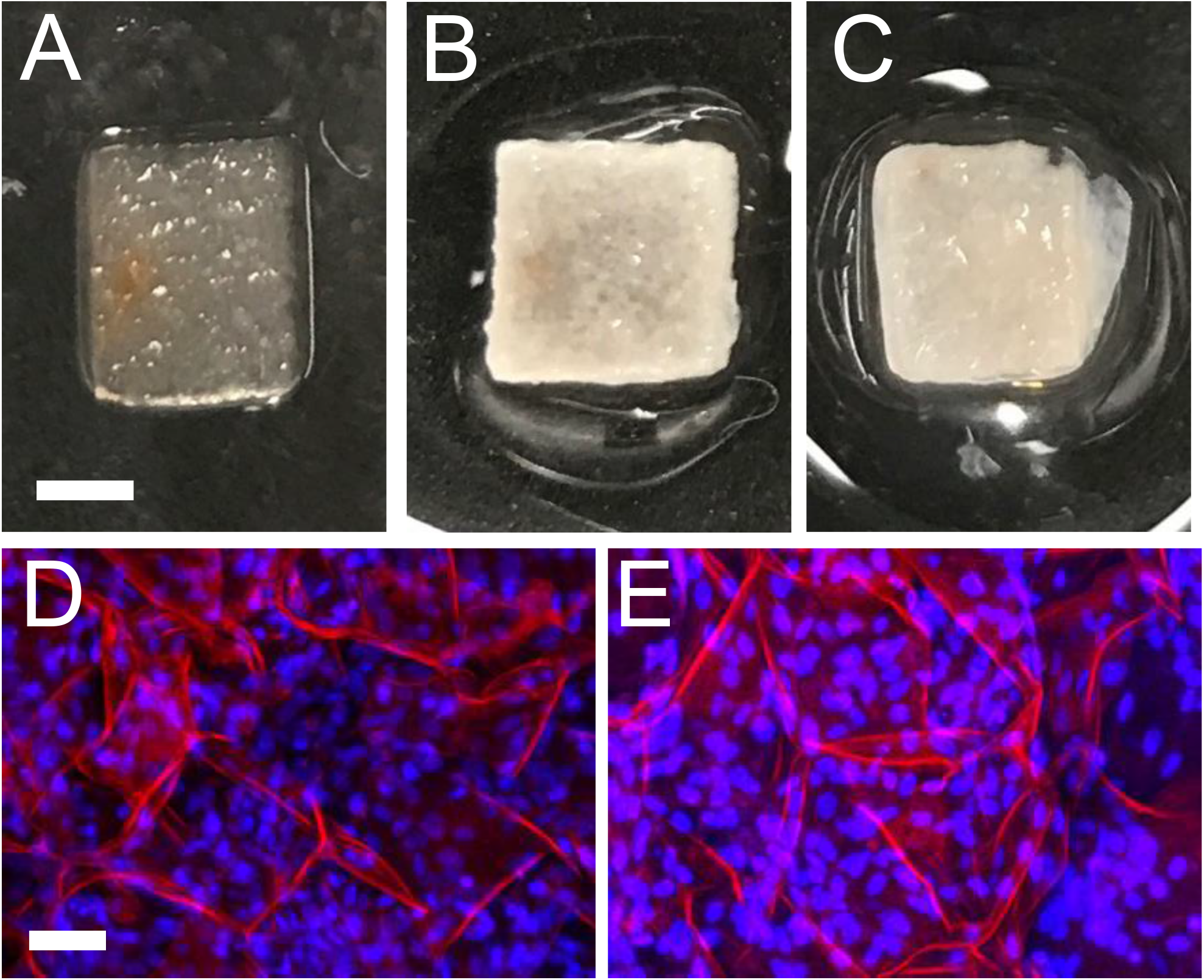
Photographs of an apple-derived cellulose scaffold after removal of the plant cells and surfactant (A) (scale bar = 2 mm - also applies to B and C), as well as a bare scaffold (B) and a calcified composite hydrogel scaffold (C) after 4-week in osteogenic differentiation medium. Representative confocal laser scanning microscope images showing seeded cells on a bare scaffold (D) (scale bar = 50 μm - also applies to E) and a composite hydrogel scaffold (E). The scaffolds were stained for cellulose (red) and for cell nuclei (blue) using propidium iodide and DAPI staining respectively. Three different scaffolds were analyzed for each experimental condition.

Confocal laser scanning microscopy showed that cells were homogenously distributed in the bare scaffolds as well as the composite hydrogel scaffolds (Figure 1D and E, respectively).

Finally, the decellularized apple-derived cellulose scaffolds (prior to collagen treatment and before MC3T3 cell seeding) displayed an average pore size of 154 ± 40 μm. The pore size distribution ranged from 73 μm to 288 μm, with the majority of the pores being between 100 and 200 μm (Figure 2).

**Figure 2:**
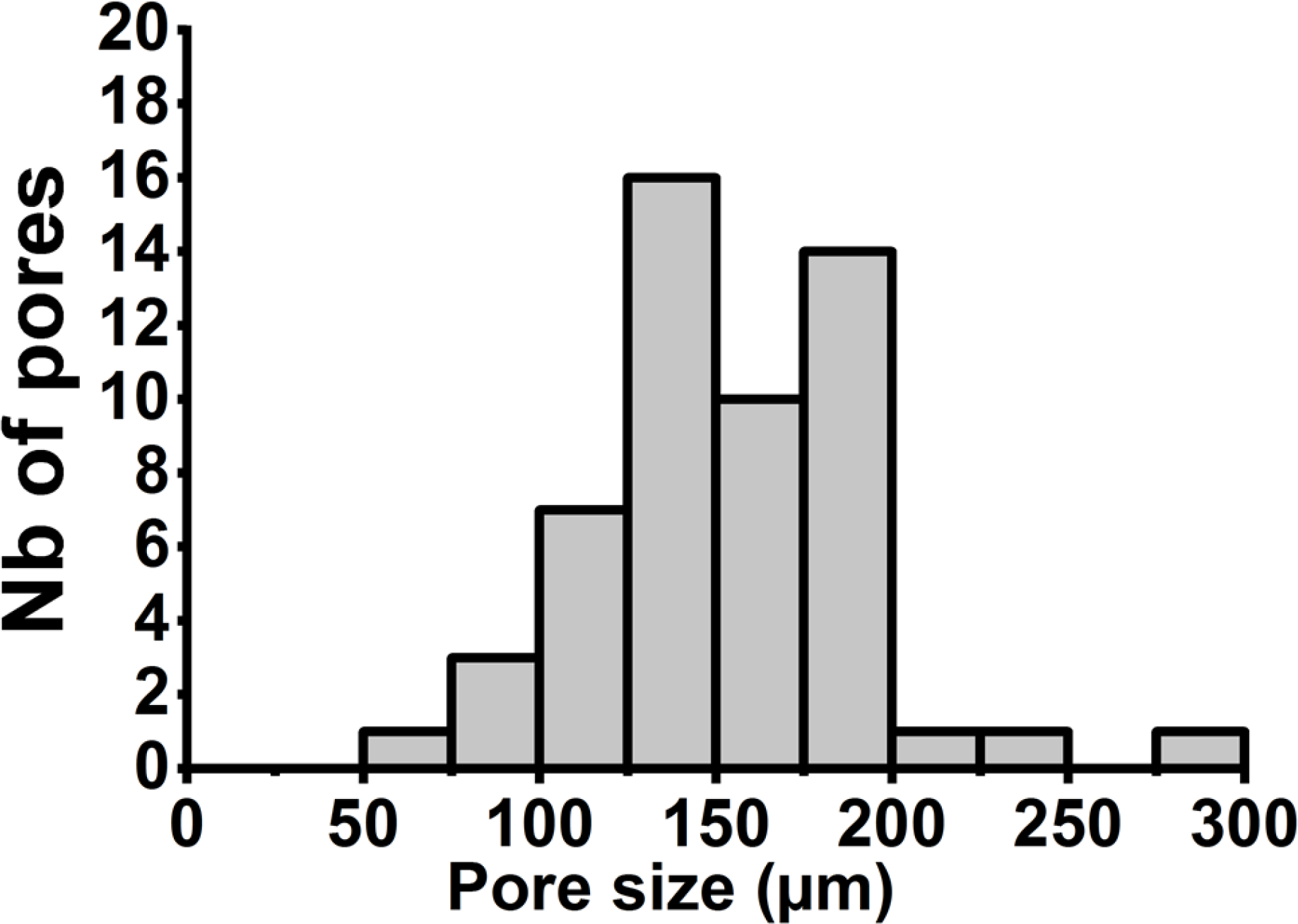
Pore size distribution of decellularized apple-derived cellulose scaffolds, before collagen treatment and MC3T3 cell seeding, from maximum projections in the Z axis of confocal images. A total of 54 pores were analyzed in 3 different scaffolds (6 pores in 3 randomly selected areas per scaffold).

### Young’s modulus measurements

The Young’s moduli of both scaffold types (bare and composite hydrogel) as well as control scaffolds (without cells) were measured after the 4 weeks of incubation in either non-differentiation or differentiation medium (Figure 3).

**Figure 3:**
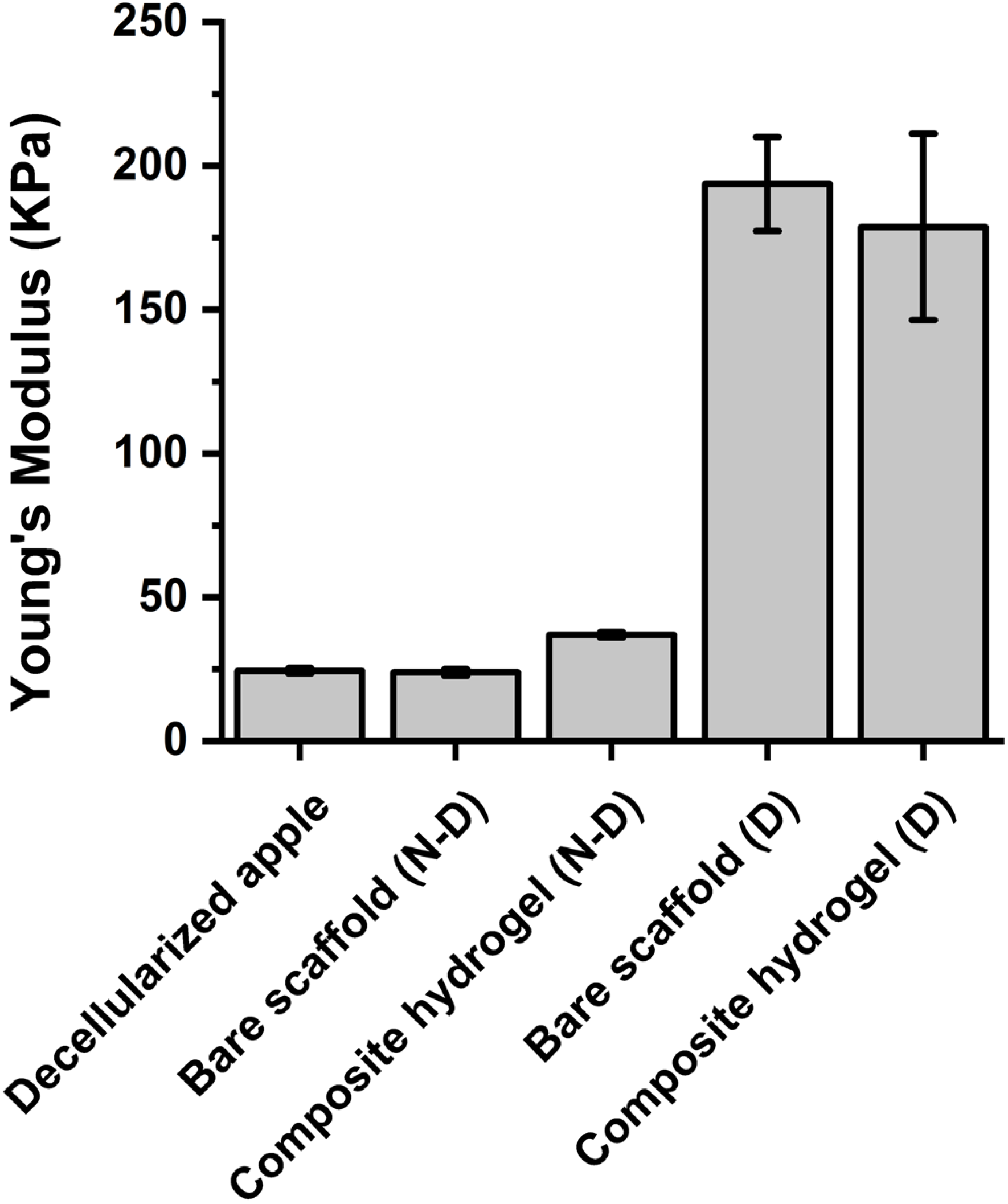
Young’s modulus of cell-seeded bare and composite hydrogel scaffolds after 4 weeks of culture in either non-differentiation or differentiation medium. Decellularized apple-derived cellulose scaffolds without cells served as a control. Statistical significance (described in the results section) was determined using a one-way ANOVA and Tukey post-hoc tests. (N-D) and (D): scaffolds incubated in non-differentiation and differentiation medium, respectively. Data are presented as means ± S.E.M. of three replicate samples per condition.

Results showed no significant difference in the Young’s modulus between the control scaffolds (scaffolds without cells) (24.4 ± 0.9 kPa) and the bare scaffolds as well as the composite hydrogel scaffolds cultured in non-differentiation medium (23.9 ± 1.2 kPa and 36.9 ± 1.0 kPa, respectively) (Figure 3). On the other hand, a significant difference was observed between the control scaffolds (24.4 ± 0.9 kPa) and the bare scaffolds as well as the composite hydrogel scaffolds cultured in differentiation medium (193.8 ± 16.4 kPa and 178.9 ± 32.4 kPa, respectively; *p*<0.001 in both cases). Furthermore, the Young’s moduli of the scaffolds cultured in non-differentiation and differentiation media were significantly different for both the bare and the composite hydrogel scaffolds (*p*<0.001 in both cases). However, there was no significant difference between the Young’s moduli of the bare and the composite hydrogel scaffolds cultured in either non-differentiation or differentiation medium.

### Alkaline phosphatase and Alizarin Red S staining

To analyze ALP activity and mineralization, the scaffolds were stained with BCIP/NBT and ARS, respectively (Figure 4).

**Figure 4:**
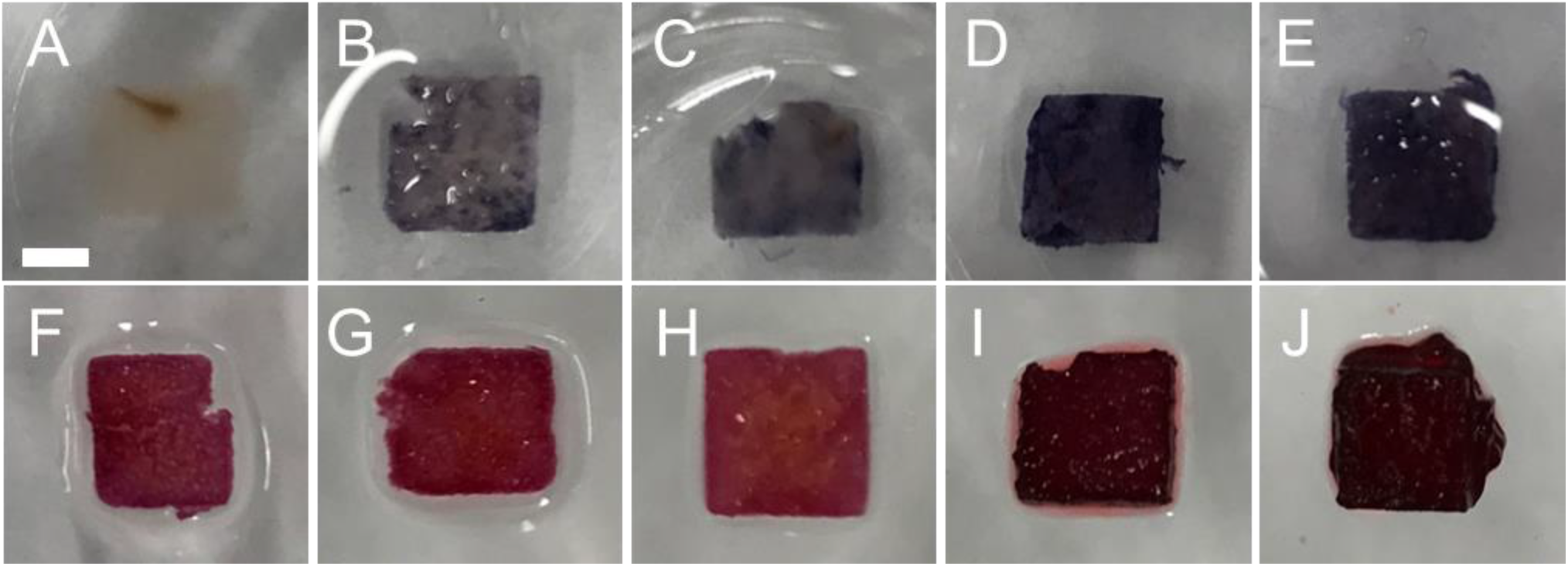
Photographs of scaffolds stained with 5-bromo-4-chloro-3’-indolyphosphate and nitro-blue tetrazolium (BCIP/NBT) (A-E) or Alizarin Red S (ARS) (F-J) (scale bar in A = 2 mm - applies to all). The alkaline phosphatase (ALP) activity was visualized using BCIP/NBT staining. The control scaffolds (bare scaffolds without cells) (A) did not stain with BCIP/NBT. Stronger ALP activity was visualized by stronger blue contrast in the bare scaffolds (D) and the composite hydrogel scaffolds (E) containing differentiated cells, compared to their counterparts with non-differentiated cells (B and C, respectively). For the ARS staining, the control scaffolds (bare scaffolds without cells) (F), the bare scaffolds with non-differentiated cells (G) and the composite hydrogel scaffolds with non-differentiated cells (H), displayed a light red color. The calcium deposition was highlighted with a strong, dark red color in the bare scaffolds (I) and the composite hydrogel scaffolds (J) containing differentiated cells. Three different scaffolds were analyzed for each experimental condition.

BCIP/NBT staining (reflecting ALP activity) was much stronger in the bare scaffolds and the composite hydrogel scaffolds with differentiated cells (Figure 4D and E, respectively) than in the scaffolds (both types) with non-differentiated cells (Figure 4B and C, respectively). The control scaffolds (scaffolds without cells) did not show any staining (Figure 4A). In addition, no difference in staining was observed between the bare scaffolds and the composite hydrogel scaffolds cultured in either non-differentiation (Figure 4B and C) or differentiation medium (Figure 4D and E).

Cells in both the bare and the composite hydrogel scaffolds cultured in differentiation medium displayed a stronger red color after ARS staining (Figure 4I, J) than cells in the scaffolds (both types) cultured in non-differentiation medium (Figures 4G, H). Control scaffolds (without cells) and scaffolds with cells cultured in non-differentiation medium displayed a non-specific staining, but this coloration was much lighter (Figure 4F-H).

### Histological analysis

Histological analysis was used to evaluate cell infiltration and scaffold mineralization. The scaffolds were fixed, embedded in paraffin, and stained with H&E or VK. Cell infiltration was demonstrated using H&E (Figure 5A, B, E, F) and scaffold mineralization was analyzed using VK staining (Figure 5C, D, G, H)

**Figure 5:**
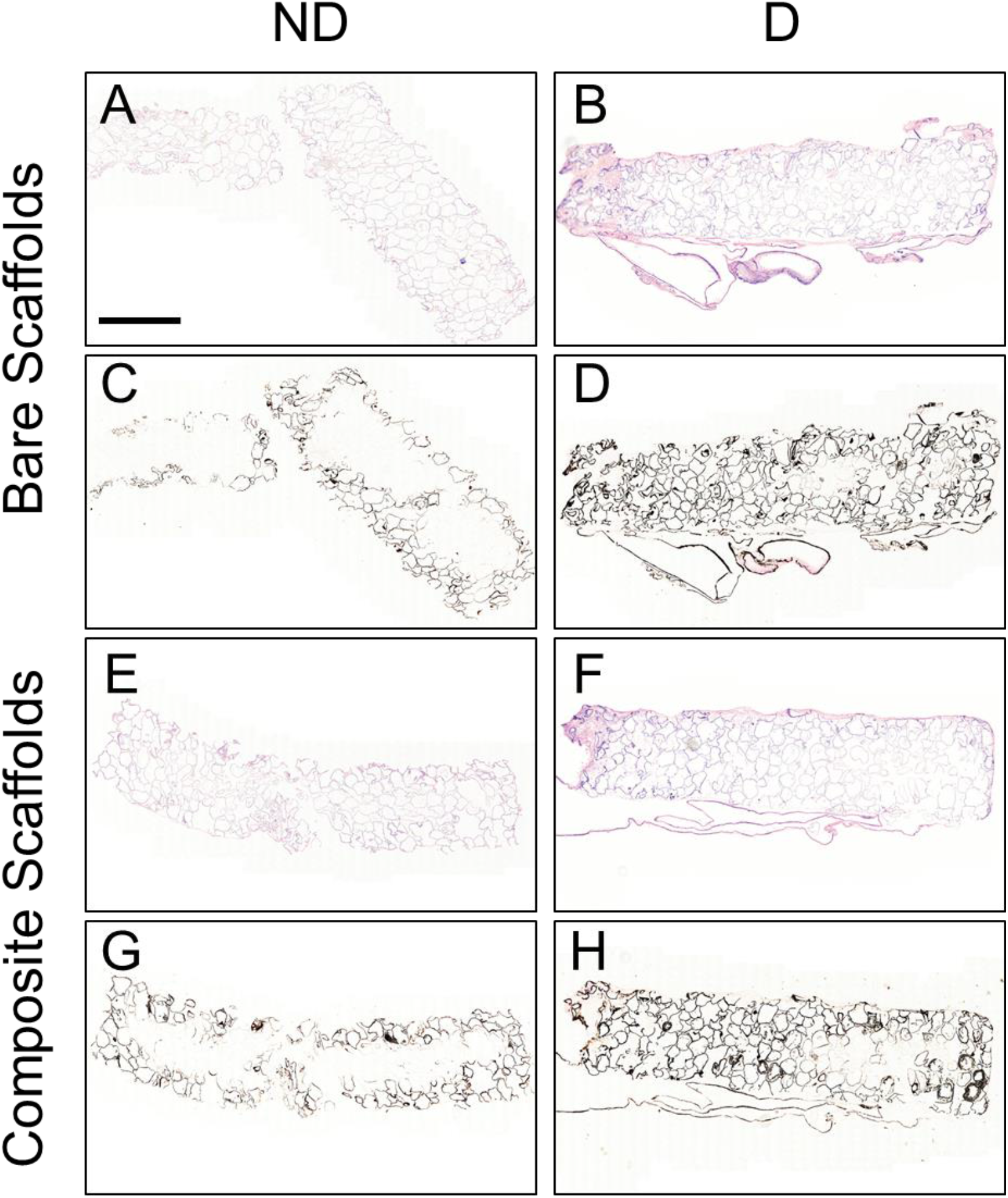
Representative images of scaffold histological cross-sections. Paraffin-embedded scaffolds were cut into 5 μm-thick sections and stained with Hematoxylin and Eosin (H&E) to visualize cell invasion (A, B, E and F) or Von Kossa (VK) to visualize mineralization (C, D, G and H) (scale bar in A = 1 mm - applies to all). Bare scaffolds and composite hydrogel scaffolds were infiltrated with MC3T3-E1 cells with multiple nuclei and cytoplasm visible at the periphery and throughout the scaffolds (A, B, E and F, blue and pink, respectively). Collagen was also visible in pale pink and more pronounced in the composite hydrogel scaffolds. The pore walls in the bare scaffolds and in the composite hydrogel scaffolds only showed the presence of mineralization at the periphery of the scaffolds when cultured in non-differentiation medium (C, G). The pore walls in the bare scaffolds and in the composite hydrogel scaffolds were entirely stained in black when cultured in differentiation medium (D, H). The bare scaffolds cultured in non-differentiation medium were damaged upon sectioning (A, C). (N-D) and (D): scaffolds incubated in non-differentiation and differentiation medium, respectively. The analysis was performed on one scaffold of each type cultured in non-differentiation medium and on 2 scaffolds of each type cultured in differentiation medium.

Bare scaffolds and composite hydrogel scaffolds were completely infiltrated with MC3T3-E1 cells (Figure 5). Multiple nuclei and cytoplasm were visible in the periphery and through the constructs (Figure 5 A, B, E, F, blue and pink, respectively). Collagen was also visible in pale pink and more pronounced in the composite hydrogel scaffolds. The pore walls in the bare scaffolds and composite hydrogel scaffolds were entirely stained in black after the 4-weeks of culture in differentiation medium (Figure 5G and H, respectively). The pore walls of the bare scaffolds and the composite hydrogel scaffolds cultured in non-differentiation medium only showed the presence of mineralization on the outside periphery of the constructs (Figure 5C and D, respectively).

### Mineralization analysis using scanning electron microscopy and energy-dispersive spectroscopy

Samples were fixed and imaged using SEM for mineral aggregates. EDS was performed to analyze the chemical composition of the aggregates.

Localized mineralization was visible in the bare scaffolds and the composite hydrogel scaffolds seeded with cells after 4 weeks of culture in differentiation medium (Figure 6A and B, respectively). Mineral deposits appeared as globular aggregates on the edge of the pores for both types of scaffolds. No mineral aggregates were visible on the bare scaffolds without cells (Figure 6C). EDS spectra were acquired on selected regions of interest, namely on the mineral aggerates for the cell-seeded scaffolds (Figure 6D and E) and on pore walls for the non-seeded scaffolds used as a control (Figure 6F). The spectra displayed stronger signal of phosphorous (P) and calcium (Ca) in both types of scaffolds cultured in differentiation medium, compared to the non-seeded scaffolds.

**Figure 6:**
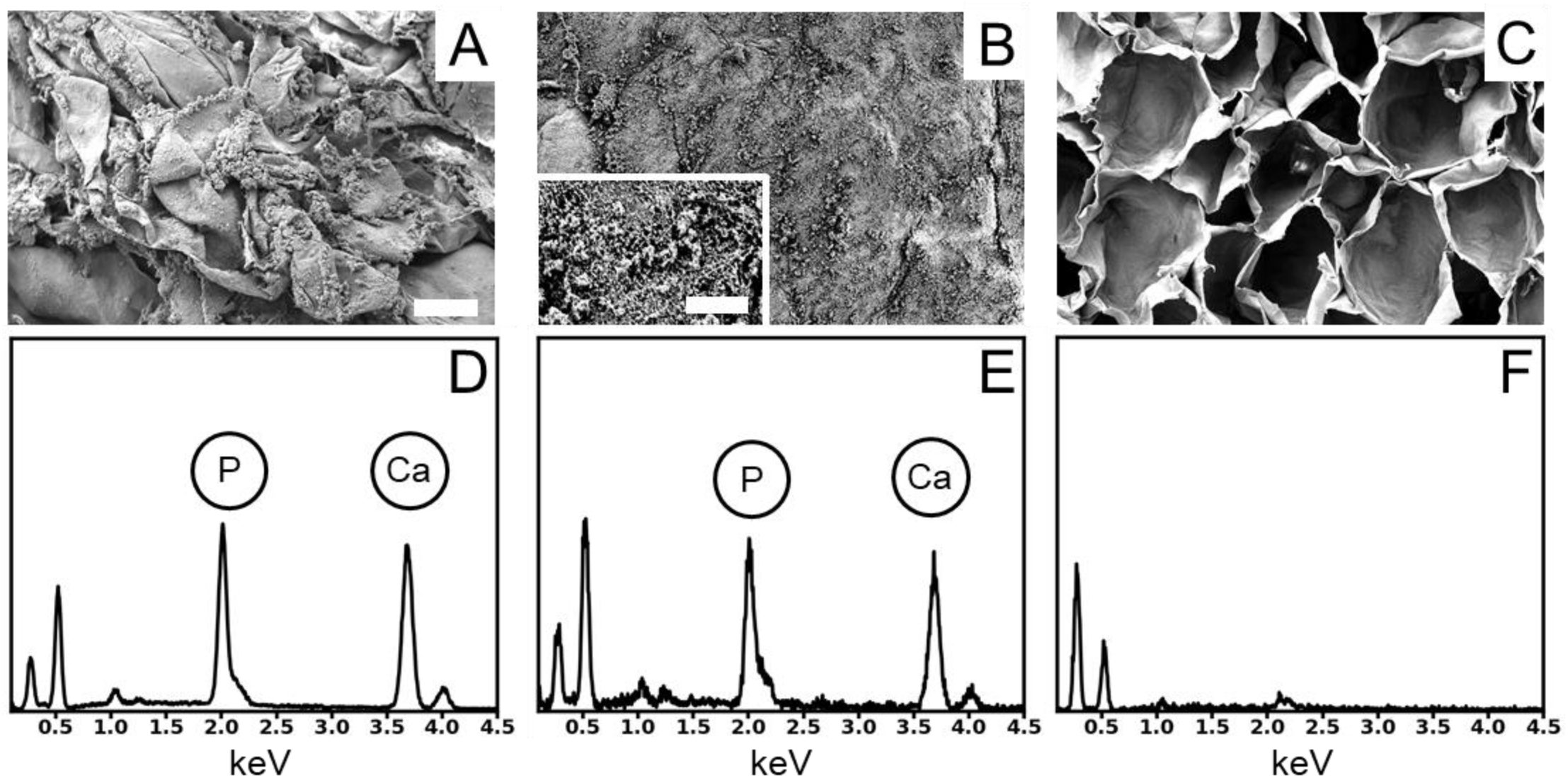
Representative scanning electron microscopy micrographs (A-C) and energy-dispersive spectra (D-F): Bare scaffold (A) and composite hydrogel scaffold (B) with MC3T3-E1 cells after differentiation, along with non-seeded cellulose scaffold (C), were gold-coated and imaged using a JEOL JSM-7500F FESEM scanning electron microscope at 2.0 kV (scale bar in A = 20 μm - applies to all). Collagen fibres are visible (B inset, scale bar = 3 μm). Energy-dispersive spectroscopy spectra were acquired on aggregates on each scaffold. Phosphorus (2.013 keV) and calcium (3.69 keV) peaks are indicated on each spectrum. Three different scaffolds were analyzed for each experimental condition.

## Discussion

Plant-derived cellulose biomaterials have potential in various fields of regenerative medicine. *In vitro* and *in vivo* studies have shown the biocompatibility of plant-derived cellulose and their potential use for tissue engineering [14]–[18]. The aim of this study was to demonstrate the potential of plant-derived cellulose to be used as a material for BTE. After removing the native cells from the apple tissue, pre-osteoblast cells (MC3T3-E1) were seeded in either bare scaffolds or composite hydrogel scaffolds (scaffolds coated with a collagen solution). The cells were let to proliferate and infiltrate the scaffold constructs for 14 days before inducing osteogenic differentiation by using differentiation medium for 4 weeks (scaffolds cultured in non-differentiation medium served as a control).

Using confocal microscopy, compression measurements, mineralization staining, histology, SEM and EDS, we confirmed that the cells were able to proliferate and differentiate within the scaffolds, thereby demonstrating the potential for plant-derived cellulose scaffolds to support bone formation. Confocal laser scanning microscopy confirmed that the cells adhered to the bare cellulose scaffolds and the composite hydrogel scaffolds (Figures 1D and E, respectively). Interestingly, calcium deposits were observed in the scaffolds (Figures 1B and C), and more specifically on the edge of the pores. The shape (globular) of these aggregates for both types of scaffolds was similar to the one reported in a previous study [21]. In addition, a large number of cell nuclei was observed around the cellulose pores as well as inside the scaffold pores (Figures 1D and E). This is consistent with previous studies which showed that various types of cells successfully adhered and proliferated onto plant-derived cellulose scaffolds [14], [16]. Moreover, similarly to one of our previous study [14], we observed that the diameter of the scaffold individual pores was about 154 μm, with the majority of the pores being between 100 and 200 μm (Figure 2). This is in line with the optimum pore size for bone growth, which has been shown to be in the range of 100–200 μm [7].

Furthermore, a significant change (about 3 to 8-fold increase) in the Young’s modulus of both the bare scaffolds and the composite hydrogel scaffolds was demonstrated after culture in differentiation medium (Figure 3). On the other hand, the addition of cells in the bare or the composite hydrogel scaffolds cultured in non-differentiation medium did not significantly affect the Young’s modulus of the constructs, and the modulus was similar to that of the control scaffolds (without cells). Interestingly, no significant differences were observed between the bare scaffolds and the composite hydrogel scaffolds cultured in either non-differentiation or differentiation medium. Overall, these results indicate that the mineralization in either type of scaffolds cultured in differentiation medium resulted in an increase of the Young’s modulus, but the presence of type 1 collagen gel in the composite scaffolds did not further increase the Young’s modulus. It should be noted that despite the increase in the Young’s modulus of both types of scaffolds when cultured in differentiation medium, the moduli remained much lower than that of bone (0.1 to 2 GPa for trabecular bone and 15 to 20 GPa for cortical bone [8]). These scaffolds may therefore not be appropriate for load-bearing applications, but still remain promising for non-load bearing applications (e.g., fractures in hand and wrist).

Staining results revealed a higher expression of ALP (Figure 4D and E) and the presence of more calcium deposits (Figure 4D and E) within both types of scaffolds after 4 weeks of culture in differentiation medium (Figure 4I and J) than in the control scaffolds (Figure 4A and F) and in both types of scaffolds cultured in non-differentiation medium (Figure 4B, C and G, H, respectively). Histological analysis showed invasion and proliferation of MC3T3-E1 cells in both types of scaffolds (Figure 5A, B, E, F), with also a similar cell distribution. The pore walls of the constructs were mineralized by the osteoblasts after the 4-week differentiation period (Figure 5D and H) in both types of scaffolds. Of note is that the periphery of the constructs with non-differentiated cells was also stained with VK. This non-specific staining may have been due to residual CaCl_2_ in the scaffolds after the decellularization process. Visual confirmation of mineralization was further assessed by qualitative analyses of SEM pictures. After the 4-week period in differentiation medium, both cell-seeded scaffold types displayed signs of ECM mineralization. Indeed, aggregates of minerals were visible on the scaffold constructs, specifically on the edges of the pores, which corroborate a previous study by Addison et al. [22] using MC3T3-E1 extracellular matrix. These aggregates were not visible on the bare scaffolds without cells. EDS analysis of the aggregates revealed high level of P and Ca, thereby suggesting the presence of apatite on the scaffold constructs.

## Conclusion

This study showed that pre-osteoblasts can adhere and proliferate within apple-derived cellulose scaffold constructs, either untreated or coated with a collagen solution. Mineralization occurred within both types of scaffolds after chemically inducing osteogenic differentiation of pre-seeded pre-osteoblasts, which resulted in an increase in the Young’s modulus of the constructs. Interestingly, these apple-derived scaffolds had a suitable pore size for BTE applications. Overall, results reveal the potential of plant-derived cellulose as a possible scaffold candidate for BTE applications. Further experiments, such as critical-size defect regeneration in animal models, are essential to confirm the applicability of this biomaterial candidate for BTE.

## Acknowledgments

This work was supported by a Natural Sciences and Engineering Research Council (NSERC) Discovery Grant. M.L.L. was supported by the Ontario Centers of Excellence *TalentEdge* program. R.J.H. was supported by an NSERC postgraduate scholarship and an Ontario Graduate Scholarship (OGS).

